# *aPEAR:* an R package for autonomous visualisation of pathway enrichment networks

**DOI:** 10.1101/2023.03.28.534514

**Authors:** Ieva Kerseviciute, Juozas Gordevicius

**Affiliations:** VUGENE, Vilnius, Lithuania

## Abstract

**Summary:** The interpretation of pathway enrichment analysis (PEA) results is frequently complicated by an overwhelming and redundant list of significantly affected pathways. Here, we present an R package *aPEAR* (Advanced Pathway Enrichment Analysis Representation) which leverages similarities between the pathway gene sets and represents them as a network of interconnected clusters. Each cluster is assigned a meaningful name which highlights the main biological themes in the experiment. Our approach enables automated and objective overview of the data without manual and time-consuming parameter tweaking.

**Availability and implementation:** The package *aPEAR* is implemented in R, published under the MIT open source licence. The source code, documentation, and usage instructions are available on https://gitlab.com/vugene/aPEAR as well as on CRAN (https://CRAN.R-project.org/package=aPEAR).

**Supplementary information:** The complete analysis used to evaluate the package can be found at https://github.com/ievaKer/aPEAR-publication.

## 1 Introduction

Pathway enrichment analysis (PEA) is indispensable when interpreting high-throughput omics data and identifying the underlying biological processes that are dysregulated in a particular condition or disease (Khatri, Sirota and Butte 2012). Despite the comprehensive insights provided by the vast number of available pathway gene set annotations in various databases, analysing large amounts of pathways introduces the redundancy problem: a single gene can be involved in multiple biological processes, resulting in pathways being highly correlated and containing overlapping sets of genes (Merico et al. 2010, Reimand et al. 2019). This causes a profusion of significantly affected pathways and impedes the interpretation of the PEA results. Ultimately, there is a need to aggregate similar pathways and analyse their interactions.

Here, we present an R package *aPEAR* (Advanced Pathway Enrichment Analysis Representation), which aids in the interpretation of the PEA results. The *aPEAR* package implements multiple metrics to calculate similarities between pathway gene sets, detects pathway clusters, and assigns biologically relevant names to them. Finally, *aPEAR* builds a visual representation of an enrichment network that can be explored interactively to elucidate the biological processes affected between the experimental conditions.

## 2 Methods

The R package *aPEAR* exports a single main function *enrichmentNetwork()* which visualises the PEA results as a network where nodes and edges represent the pathways and similarity between them, respectively. While it was created with the clusterProfiler (Wu et al. 2021) and gprofiler2 (Kolberg et al. 2020) output in mind, any enrichment result is accepted as long as it is formatted correctly (Supplementary Text 1). The network is constructed in several steps:

1. The pairwise similarity between all pathway gene sets is evaluated using the Jaccard index (default), cosine similarity, or correlation similarity metrics.
2. The similarity matrix is then used to detect clusters of redundant pathways using Markov (default) (Van Dongen 2008), hierarchical, or spectral (John et al. 2020) clustering algorithms.
3. Each cluster is assigned a biologically meaningful name. Network analysis is used to determine the pathway with the most connections, using either PageRank (Page et al. 1999) (default) or HITS (Kleinberg 1999) algorithm. Alternatively, the highest absolute NES value or the lowest p-value can be used to select the most important pathway in the cluster. The description of this pathway is used as the cluster label.
4. A *ggplot2* (Wickham 2016) graph is constructed using the similarity matrix and the annotated clusters. An interactive graph is visualised using *plotly* (Sievert 2020). Pathways and their assigned clusters are returned as output as well (Supplementary Text 2).

## 3 Results

Currently, the most frequently used tools for gene set visualisation include the *emapplot* function from the R package *enrichplot* (Yu 2022) and the Cytoscape plugin *Enrichment Map* (Merico et al. 2010). These methods use a word cloud-like algorithm to assign a cluster name which results in labels that are not semantically meaningful. The *emapplot* has some limitations when handling large datasets and produces numerous small clusters with non-intuitive labels (Supplementary Methods, Supplementary Fig. 1). Cytoscape, while a powerful software tool that is able to work with large amounts of data, requires many time consuming manual adjustments that can introduce bias into the network (Supplementary Methods, Supplementary Fig. 2, Supplementary Fig. 3, Supplementary Fig. 4). In contrast, *aPEAR* makes it easy to work with numerous pathways, highlights the biological context of the clusters and does not require additional manipulation of the graph, making it ideal for automated data visualisation as well as interactive investigations (Fig. 1).

**Fig. 1.**
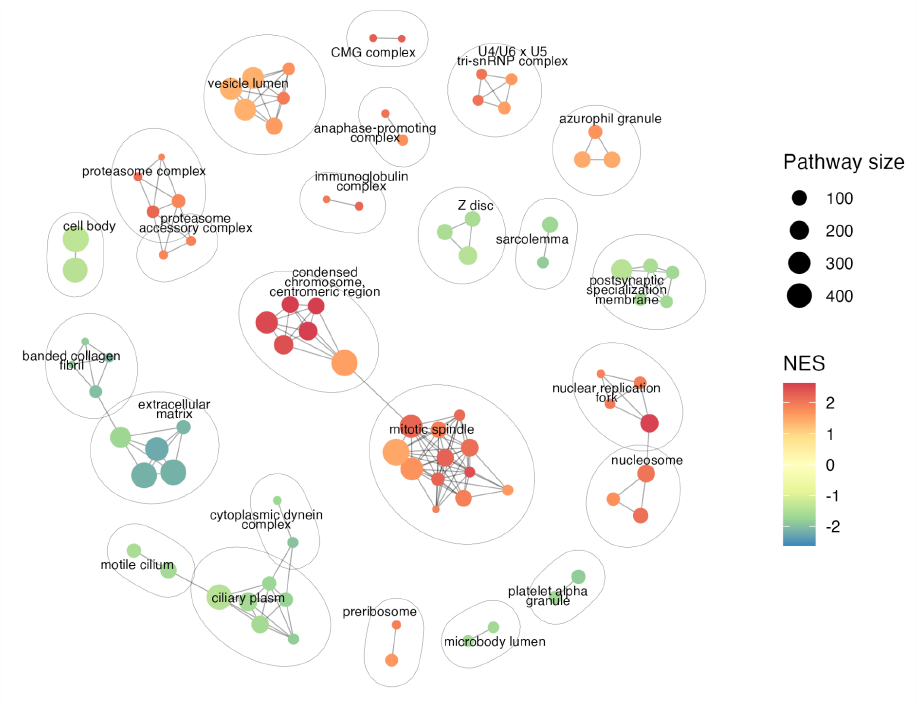
Example enrichment network generated by *aPEAR*. The nodes represent the significant pathways and the edges represent similarity between them. Coloured by normalised enrichment score (NES).

To determine which similarity metric and clustering algorithm is best suitable for pathway cluster analysis, 180 tests were performed using PEA results from 10 real-world datasets (Supplementary Methods, Supplementary Table 1, Supplementary Table 2). Based on cluster quality evaluation using the Dunn index (Dunn 1974), the Silhouette index (Rousseeuw 1987) and the Davies-Bouldin index (Davies and Bouldin 1979), the Jaccard similarity metric and the Markov clustering algorithm were found to be best suited for such analysis and, thus, were set as the default parameters in the *aPEAR* package (Supplementary Text 3, Supplementary Fig. 5). Note that the Jaccard coefficient can be affected by the pathway size and may under-connect the smaller pathways contained within the larger ones (Salvatore et al. 2020).

## Conclusion

We developed an R package–*aPEAR*–that visualises clusters of similar pathways as an enrichment network and, consequently, enables better interpretation of the pathway enrichment analysis results.

## Acknowledgements

We would like to express our sincere gratitude to Migle Gabrielaite for her invaluable manuscript reviews and to Milda Milčiūtė for her excellent work in testing the *aPEAR* package.

