## Supplementary file for "*aPEAR:* an R package for autonomous visualisation of pathway enrichment networks"

### Supplementary information

**Supplementary Text 1.** The detailed description of the input format for the *aPEAR* package.

***For more detailed and up-to-date information, please check the vignette of the aPEAR package.***

The most important function exported by the *aPEAR* package is `enrichmentNetwork()`. This function accepts the enrichment results as a *data.frame* object. When using *gprofiler2* (Kolberg *et al.* 2020) or *clusterProfiler* (Wu *et al.* 2021), the *aPEAR* package will handle all necessary actions when provided with the *data.frame* of the results from these methods, for example:

```
> # Perform enrichment analysis
> enrich <- clusterProfiler::gseGO(geneList, OrgDb = org.Hs.eg.db, ont = "CC")
>
> # Generate enrichment network using aPEAR
> aPEAR::enrichmentNetwork(enrich@result)
```

When using gprofiler2, do not forget to set `evcodes = TRUE` in order to obtain the gene list for each pathway.
In case the enrichment analysis is carried out differently, the package can still be utilised to create the enrichment network. The data should be formatted as a *data.frame* object that comprises the following columns:

- ***Description*** - this column contains the titles of the pathways and will be used to generate the cluster names.
- ***pathwayGenes*** - this column contains the list of genes in the pathway. The user may choose to use the complete list of genes in the pathway or only the core enrichment genes.
- A column that will be used to colour the nodes (we recommend using p-values or NES). The user has to specify which column to use with `enrichmentNetwork(..., colorBy = "columnName")`.
- A column that will be used to choose the size of the node (we recommend using the number of genes in the pathway). The user has to specify which column to use with `enrichmentNetwork(..., nodeSize = "columnName")`.

The ***Description*** and ***pathwayGenes*** columns for custom inputs are **required**. The columns for node size and colour have to be specified, so the user can choose them freely. For example:

```
> # This is an example of correctly formatted enrichment result
> # (can contain other columns as well):
> enrich[ 1:5, ]

##                               Description                          pathwayGenes       NES size
## 1: Abc-Family Proteins Mediated Transport  11160/27248/51009/6400/10956/... -2.865165  103
## 2:        Acetyl-Coa Biosynthetic Process       1738/1737/5162/38/23417/... -2.442777   13
## 3:           Acetyl-Coa Metabolic Process        1738/1737/5162/38/3158/... -2.655216   24
## 4:          Activation Of Immune Response      929/7100/1191/6850/10392/...  2.201129  300
## 5:          Acyl-Coa Biosynthetic Process 1738/1737/5162/9524/109703458/... -3.026606   38
```

```
> # Generate enrichment network using aPEAR
> aPEAR::enrichmentNetwork(enrich, nodeSize = "size", colorBy = "NES")
```

Note, that it does not matter which gene identifiers are used as long as they are the same between all pathways. For more information on the available methods and plotting parameters, please check `?aPEAR.methods` and `?aPEAR.theme`.

**Supplementary Text 2.** Usage of *aPEAR* to obtain clusters of redundant pathways.

We also provide a simple way to obtain the pathway clustering results. This can be done with `findPathClusters()`. The function accepts a *data.frame* in the same format as `enrichmentNetwork()` and returns a list of clusters (*data.frame*) and the similarity matrix (*matrix*) which can later be used to create the enrichment network by `plotPathClusters()`. These two methods are internally called by `enrichmentNetwork()` and generate the same result.

```
> # Get clustering result using aPEAR
> data <- aPEAR::findPathClusters(enrich@result)
> data$clusters[ 1:3 ]

##                                     ID                                  Cluster
## 1:                         CMG complex   DNA replication preinitiation complex
## 2:               DNA packaging complex                               nucleosome
## 3: DNA replication preinitiation complex   DNA replication preinitiation complex

> # Create the enrichment network visualisation using default parameters
> aPEAR::plotPathClusters(enrich@result, data$sim, data$clusters)
```

**Supplementary Text 3.** The evaluation of clustering quality.

To determine which similarity metric and clustering algorithm produces the best results, we performed clustering evaluation using multiple cluster quality indexes that look at various aspects of the data. Specifically, we used the Dunn index, the Silhouette index, and the Davies-Bouldin index. The Dunn index measures the separation between clusters and the compactness of each cluster (Dunn 1974). The Silhouette index indicates how well each data point fits into its assigned cluster, based on the average distance between the point and all other points in its cluster as well as the average distance between the point and all points in the nearest neighbouring cluster (Rousseeuw 1987). The Davies-Bouldin index evaluates the average similarity between each cluster and its most similar cluster (Davies and Bouldin 1979).

The clusters obtained by spectral clustering algorithm received the worst estimates by all quality indexes which significantly differed from the metrics obtained by the hierarchical and the Markov clustering algorithms ($p$-value < 0.01, Wilcoxon signed-rank test) (Supplementary Fig. 5a,c). The Dunn index estimates were similar for the hierarchical and the Markov clustering algorithms, however, the Silhouette and the Davies-Bouldin indexes showed significant preference for the Markov clustering as compared to the hierarchical clustering ($p$-value $< 0.01$, Wilcoxon signed-rank test) (Supplementary Fig. 5a,c). All cluster quality indexes showed that the similarity metric did not have an impact on the cluster quality ($p$-value $> 0.01$, Wilcoxon signed-rank test) (Supplementary Fig. 5b,c). Based on this evaluation, we decided to set the default similarity metric to the Jaccard similarity metric as it is most frequently used by other similar tools (Merico *et al.* 2010; Yu 2022), and to set the default clustering algorithm to the Markov clustering algorithm as it showed the best clustering quality. The spectral clustering did not prove to be applicable to the gene-set cluster analysis but the implementation was retained for experimental purposes, however, it should be used with care.

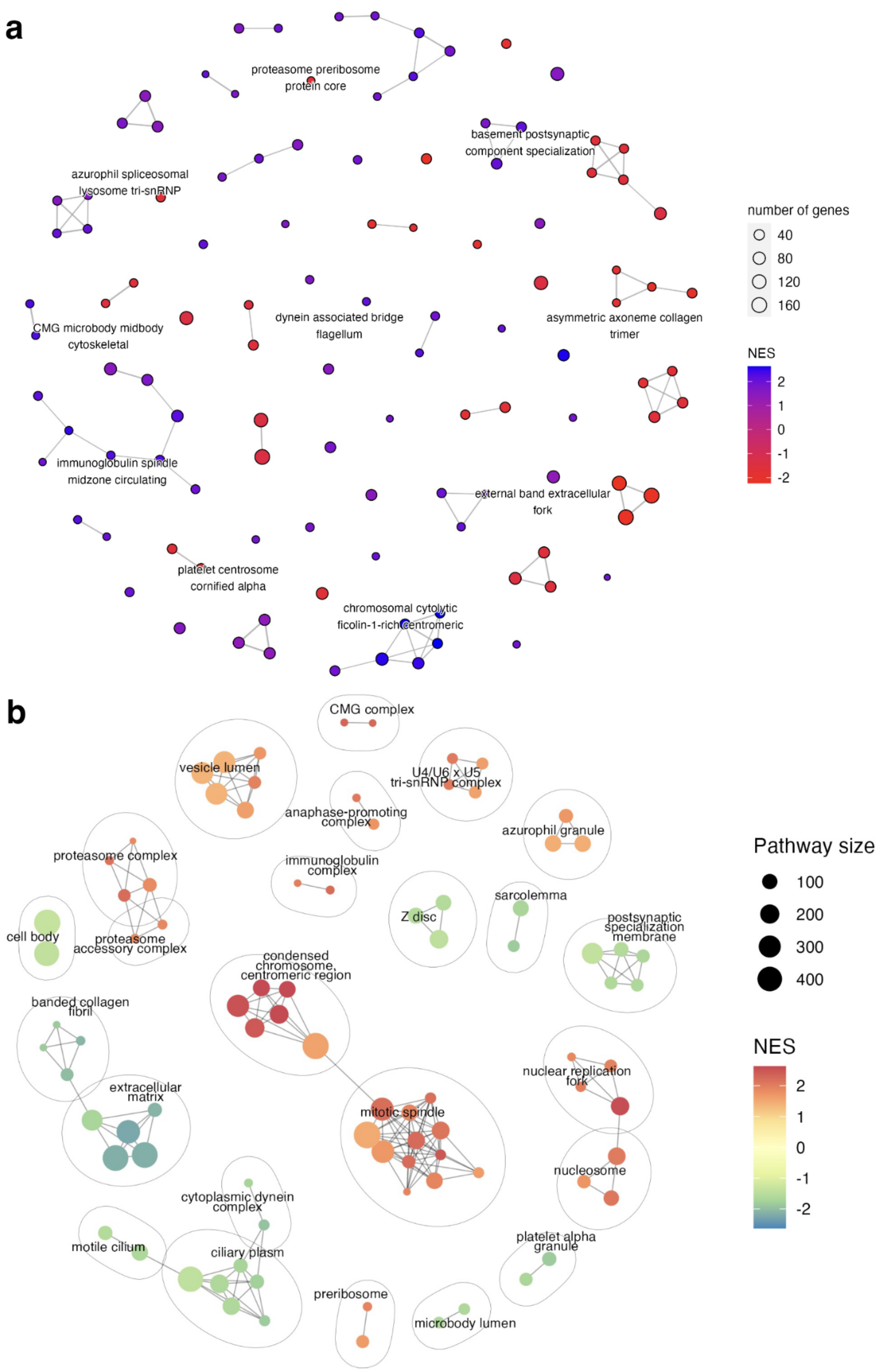


**Supplementary Figure 1.** Enrichment networks of the *clusterProfiler* results, generated using *emapplot* (a) and *aPEAR* (b). The GSEA was performed using the example gene list from the R package *DOSE*.

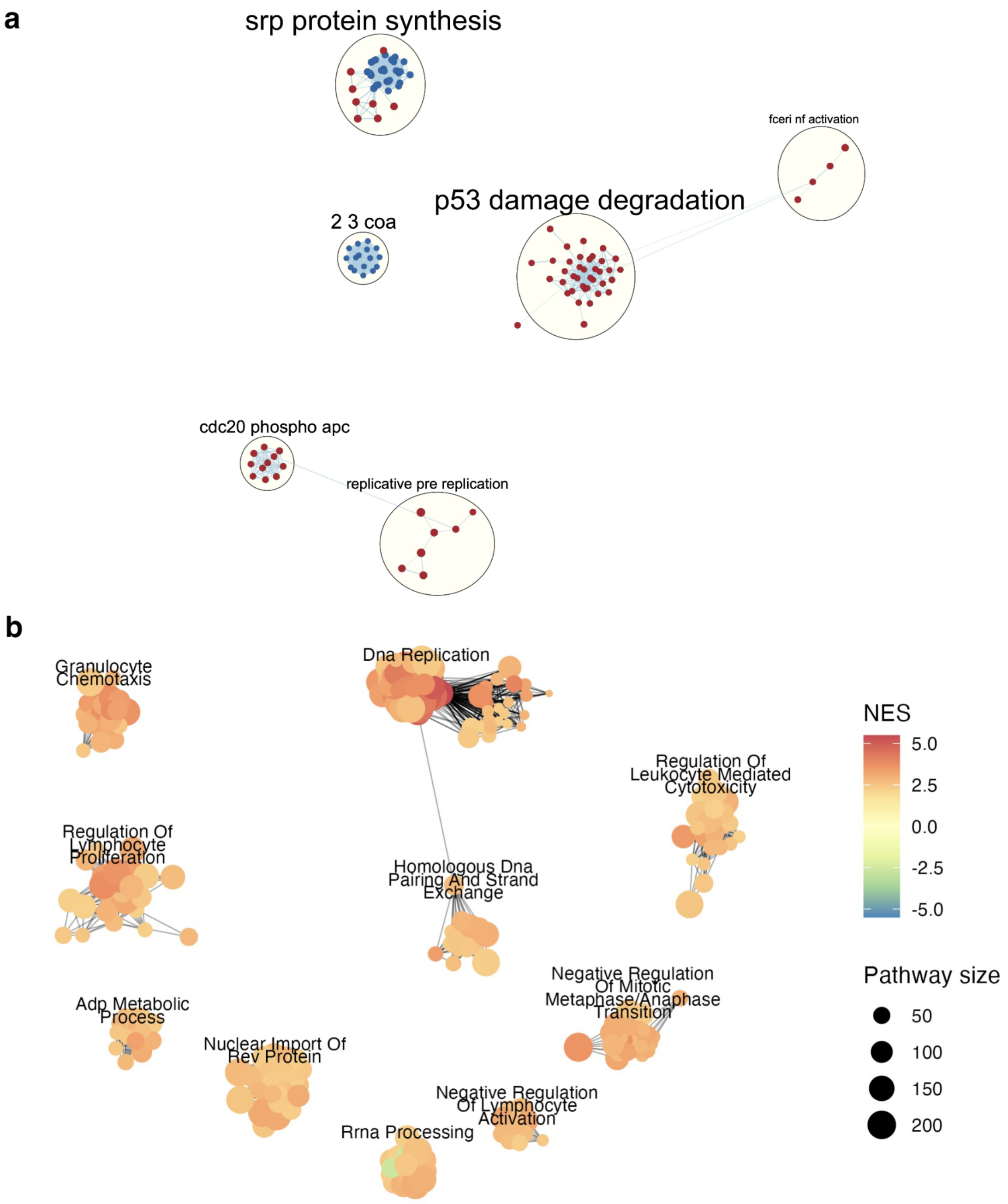


**Supplementary Figure 2.** Enrichment networks of the GSEA software results, generated using *Cytoscape* (a) and *aPEAR* (b). The GSEA was performed using the example gene list from the R package *DOSE*.

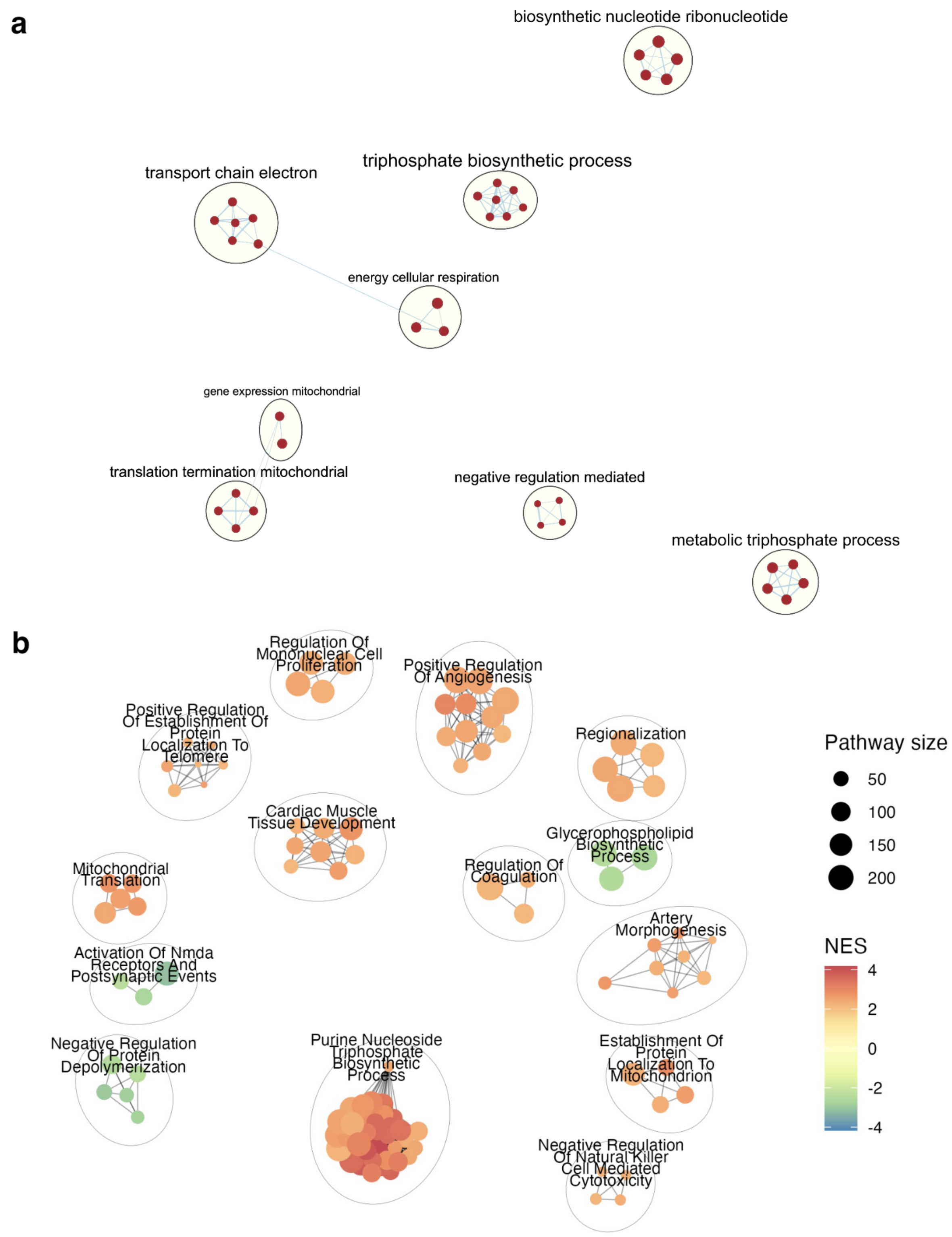


**Supplementary Figure 3.** Enrichment networks of the GSEA software results, generated using *Cytoscape* (a) and *aPEAR* (b). The GSEA was performed using a gene list from internal analysis (available for download from the GitHub repository of the publication analysis at directory *data/geneRanks/dataset2.rnk*).

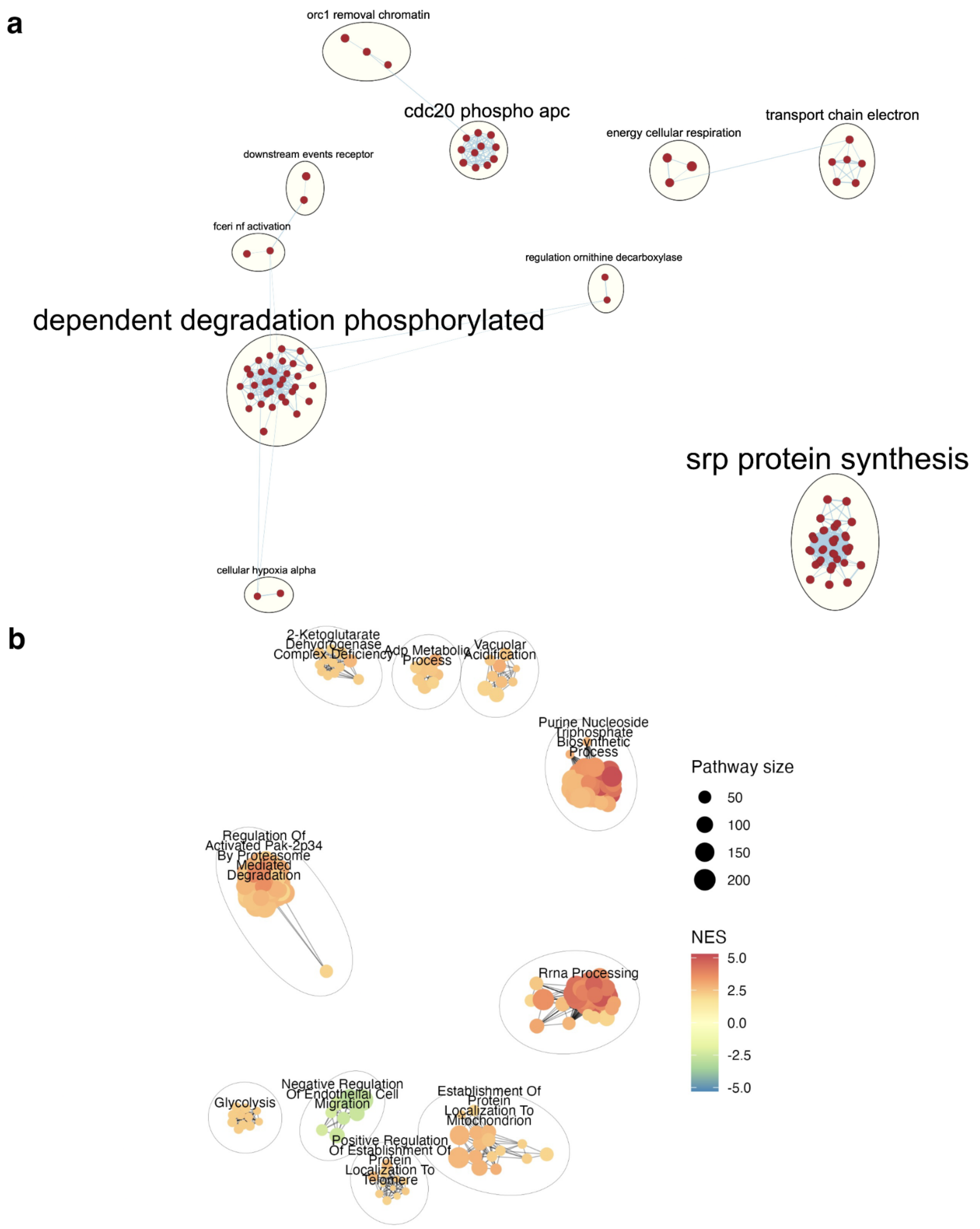


**Supplementary Figure 4.** Enrichment networks of the GSEA software results, generated using *Cytoscape* (a) and *aPEAR* (b). The GSEA was performed using a gene list from internal analysis (available for download from the GitHub repository of the publication analysis at directory *data/geneRanks/dataset3.rnk*).

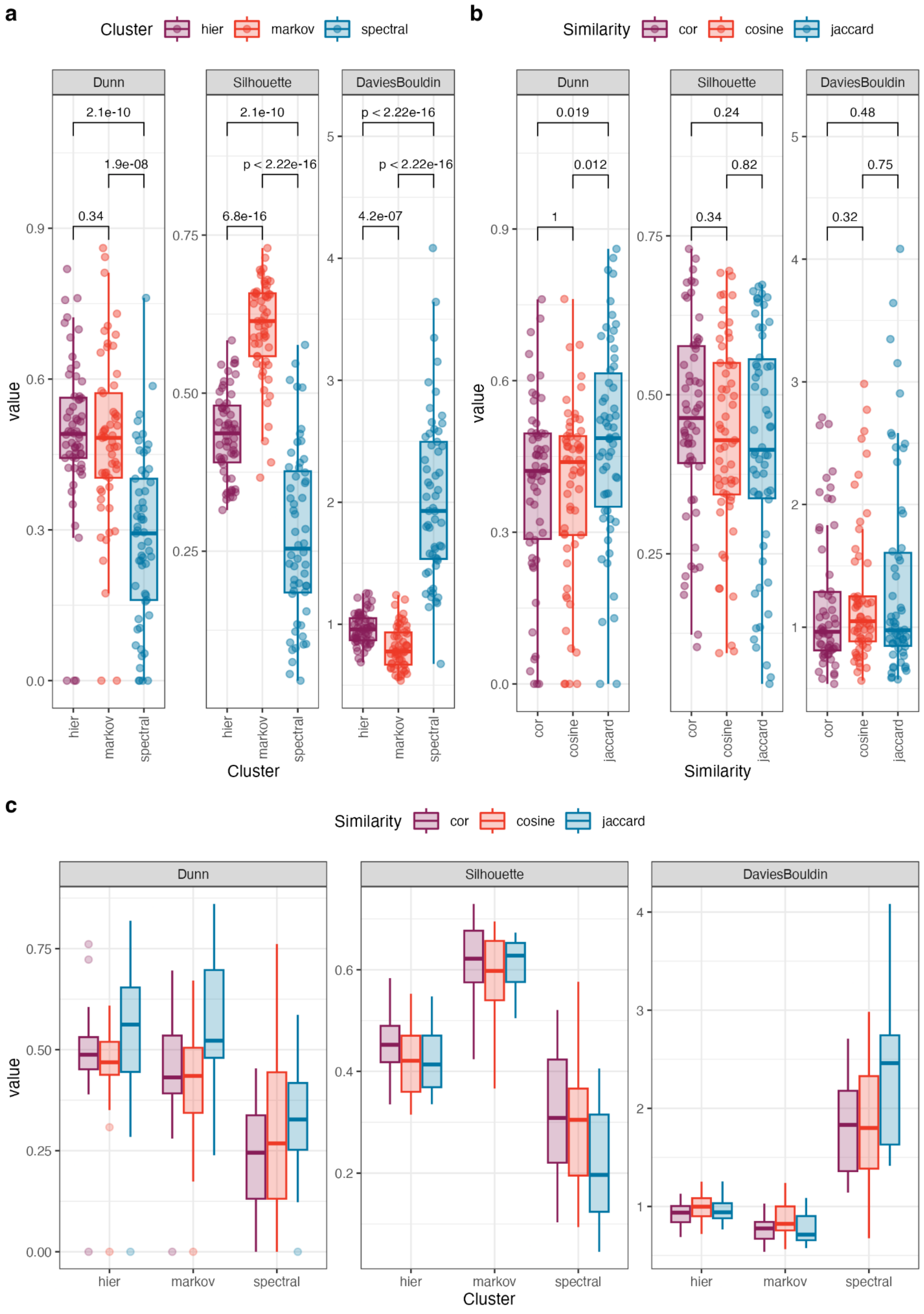


**Supplementary Figure 5.** Evaluation of cluster quality using various similarity and clustering metrics. Cluster quality dependency on clustering method (a) and similarity metric (b). The p-values obtained using the Wilcoxon signed-rank test. The figure (c) describes the cluster quality dependency on the clustering method split by the similarity metric.

**Supplementary Table 1.** Datasets used for clustering quality evaluation. 10 different real-world datasets were used to perform gene set enrichment using two methods: *gseGO* from R package *clusterProfiler* (Wu *et al.* 2021) and the GSEA software (Mootha *et al.* 2003; Subramanian *et al.* 2005). The enrichment results (top 1000 most significant pathways) can be found in the *aPEAR* publication GitHub repository.

| No | Enrichment method | Data type | Condition |
|---|---|---|---|
| 1. | *GSEA* | RNA-Seq | Alzheimer's disease |
| 2. | *clusterProfiler* | | |
| 3. | *GSEA* | RNA-Seq | SARS-CoV-2 (nasal swab) |
| 4. | *clusterProfiler* | | |
| 5. | *GSEA* | EPIC methylation | ovarian cancer |
| 6. | *clusterProfiler* | | |
| 7. | *GSEA* | EPIC methylation | persons who gave birth to newborns with FGR |
| 8. | *clusterProfiler* | | |
| 9. | *GSEA* | EPIC methylation | Alzheimer's disease |
| 10. | *clusterProfiler* | | |
| 11. | *GSEA* | EPIC methylation | mild Alzheimer's disease |
| 12. | *clusterProfiler* | | |
| 13. | *GSEA* | WGBS | severe SARS-CoV-2 |
| 14. | *clusterProfiler* | | |
| 15. | *GSEA* | EPIC methylation | persons who gave birth to newborns with autism spectrum disorder |
| 16. | *clusterProfiler* | | |
| 17. | *GSEA* | RNA-Seq | late-stage Parkinson's disease |
| 18. | *clusterProfiler* | | |
| 19. | *GSEA* | deep RNAseq | Parkinson's disease |
| 20. | *clusterProfiler* | | |

**Supplementary Table 2.** Clustering quality metrics when using various clustering and similarity metrics. Obtained using the top 100 significant pathways from the datasets described in Supplementary Table 1.

| Dataset | Cluster | Similarity | Dunn | Silhouette | DaviesBouldin |
|---|---|---|---|---|---|
| 1 | markov | jaccard | 0.23861 | 0.60606 | 0.79508 |
| 1 | hier | jaccard | 0.40959 | 0.44477 | 0.86839 |
| 1 | spectral | jaccard | 0 | 0.26236 | 1.61759 |
| 1 | markov | cosine | 0.41301 | 0.59782 | 0.75348 |
| 1 | hier | cosine | 0.37603 | 0.51779 | 0.85383 |
| 1 | spectral | cosine | 0 | 0.19516 | 2.41451 |
| 1 | markov | cor | 0.42954 | 0.57805 | 0.75033 |
| 1 | hier | cor | 0.38926 | 0.51875 | 0.86575 |
| 1 | spectral | cor | 0.05387 | 0.39577 | 1.22259 |
| 2 | markov | jaccard | 0.6635 | 0.65121 | 0.59279 |
| 2 | hier | jaccard | 0.56549 | 0.41744 | 0.92327 |
| 2 | spectral | jaccard | 0.37688 | 0.18737 | 2.34632 |
| 2 | markov | cosine | 0.5091 | 0.60969 | 0.70699 |
| 2 | hier | cosine | 0.44295 | 0.45271 | 0.83126 |
| 2 | spectral | cosine | 0.26803 | 0.24377 | 2.24216 |
| 2 | markov | cor | 0.48783 | 0.65272 | 0.62222 |
| 2 | hier | cor | 0.49179 | 0.47626 | 0.80844 |
| 2 | spectral | cor | 0.32056 | 0.33427 | 2.13894 |
| 3 | markov | jaccard | 0.37606 | 0.6521 | 0.77625 |
| 3 | hier | jaccard | 0.28424 | 0.46867 | 0.92932 |
| 3 | spectral | jaccard | 0.37467 | 0.34957 | 1.41494 |
| 3 | markov | cosine | 0.34306 | 0.69495 | 0.791 |
| 3 | hier | cosine | 0.48927 | 0.46569 | 0.98048 |
| 3 | spectral | cosine | 0.23861 | 0.36826 | 1.24758 |
| 3 | markov | cor | 0.28467 | 0.67702 | 0.63603 |
| 3 | hier | cor | 0.49577 | 0.47321 | 0.92944 |
| 3 | spectral | cor | 0.23869 | 0.43584 | 1.14038 |
| 4 | markov | jaccard | 0.52219 | 0.54446 | 0.71245 |
| 4 | hier | jaccard | 0.75718 | 0.40504 | 0.86847 |
| 4 | spectral | jaccard | 0.34263 | 0.11515 | 2.9056 |

| | | | | | |
|---|---|---|---|---|---|
| 4 | markov | cosine | 0.48289 | 0.61397 | 0.6681 |
| 4 | hier | cosine | 0.46457 | 0.41829 | 0.90512 |
| 4 | spectral | cosine | 0.06213 | 0.18261 | 2.16207 |
| 4 | markov | cor | 0.57633 | 0.6559 | 0.57983 |
| 4 | hier | cor | 0.4628 | 0.42335 | 0.85872 |
| 4 | spectral | cor | 0.24525 | 0.19943 | 2.27021 |
| 5 | markov | jaccard | 0.52629 | 0.62784 | 0.90372 |
| 5 | hier | jaccard | 0.59544 | 0.51265 | 0.84139 |
| 5 | spectral | jaccard | 0.12248 | 0.28061 | 1.92639 |
| 5 | markov | cosine | 0.29438 | 0.58975 | 0.98917 |
| 5 | hier | cosine | 0.46908 | 0.42855 | 1.01535 |
| 5 | spectral | cosine | 0.48827 | 0.36048 | 1.52205 |
| 5 | markov | cor | 0.49789 | 0.67938 | 0.76904 |
| 5 | hier | cor | 0.4831 | 0.44047 | 0.97001 |
| 5 | spectral | cor | 0.421 | 0.42437 | 1.63337 |
| 6 | markov | jaccard | 0.7299 | 0.65328 | 0.64509 |
| 6 | hier | jaccard | 0.62826 | 0.34303 | 0.97042 |
| 6 | spectral | jaccard | 0.32608 | 0.05574 | 3.64232 |
| 6 | markov | cosine | 0.53443 | 0.55015 | 0.8846 |
| 6 | hier | cosine | 0.60896 | 0.36478 | 1.02807 |
| 6 | spectral | cosine | 0.18838 | 0.09375 | 2.98372 |
| 6 | markov | cor | 0.57196 | 0.5876 | 0.82818 |
| 6 | hier | cor | 0.60547 | 0.40283 | 0.99836 |
| 6 | spectral | cor | 0.41093 | 0.10329 | 2.70963 |
| 7 | markov | jaccard | 0.45001 | 0.61459 | 0.9331 |
| 7 | hier | jaccard | 0.50947 | 0.54504 | 0.88233 |
| 7 | spectral | jaccard | 0.17282 | 0.13452 | 2.46614 |
| 7 | markov | cosine | 0.29713 | 0.68674 | 0.9232 |
| 7 | hier | cosine | 0.49631 | 0.49814 | 0.88642 |
| 7 | spectral | cosine | 0.4777 | 0.24843 | 1.92879 |
| 7 | markov | cor | 0.41317 | 0.7141 | 0.82725 |
| 7 | hier | cor | 0.56027 | 0.54695 | 0.80696 |
| 7 | spectral | cor | 0 | 0.25967 | 1.79166 |
| 8 | markov | jaccard | 0.84268 | 0.6416 | 0.58693 |

| | | | | | |
|---|---|---|---|---|---|
| 8 | hier | jaccard | 0.71178 | 0.45014 | 0.97973 |
| 8 | spectral | jaccard | 0.12994 | 0.10249 | 3.34794 |
| 8 | markov | cosine | 0.50042 | 0.65799 | 0.82203 |
| 8 | hier | cosine | 0.56195 | 0.50813 | 0.96952 |
| 8 | spectral | cosine | 0.46984 | 0.19539 | 2.53648 |
| 8 | markov | cor | 0.49589 | 0.66083 | 0.77838 |
| 8 | hier | cor | 0.72285 | 0.52989 | 0.88966 |
| 8 | spectral | cor | 0.22961 | 0.18537 | 2.64665 |
| 9 | markov | jaccard | 0.47477 | 0.50454 | 1.08605 |
| 9 | hier | jaccard | 0.55849 | 0.33547 | 1.15861 |
| 9 | spectral | jaccard | 0.32044 | 0.13277 | 2.56321 |
| 9 | markov | cosine | 0.34427 | 0.53347 | 1.20226 |
| 9 | hier | cosine | 0.46854 | 0.44189 | 1.08652 |
| 9 | spectral | cosine | 0.41842 | 0.35774 | 1.53555 |
| 9 | markov | cor | 0.414 | 0.55838 | 0.93785 |
| 9 | hier | cor | 0.4526 | 0.44345 | 1.01919 |
| 9 | spectral | cor | 0.29878 | 0.22869 | 2.0443 |
| 10 | markov | jaccard | 0.86062 | 0.60547 | 0.67671 |
| 10 | hier | jaccard | 0.81887 | 0.34703 | 1.04115 |
| 10 | spectral | jaccard | 0.51651 | 0.15485 | 2.4953 |
| 10 | markov | cosine | 0.42091 | 0.39045 | 1.05177 |
| 10 | hier | cosine | 0.47307 | 0.33934 | 1.04501 |
| 10 | spectral | cosine | 0.76148 | 0.5763 | 0.67498 |
| 10 | markov | cor | 0.46475 | 0.42401 | 0.97202 |
| 10 | hier | cor | 0.76091 | 0.39719 | 0.98471 |
| 10 | spectral | cor | 0.1604 | 0.21463 | 2.21778 |
| 11 | markov | jaccard | 0.50734 | 0.67031 | 1.00263 |
| 11 | hier | jaccard | 0.42332 | 0.44891 | 1.13631 |
| 11 | spectral | jaccard | 0.52972 | 0.23806 | 1.95272 |
| 11 | markov | cosine | 0.46571 | 0.69183 | 0.76679 |
| 11 | hier | cosine | 0.44476 | 0.3958 | 1.2173 |
| 11 | spectral | cosine | 0.39468 | 0.32456 | 1.63722 |
| 11 | markov | cor | 0.43139 | 0.69713 | 0.85612 |
| 11 | hier | cor | 0.4511 | 0.47908 | 1.09558 |

| | | | | | |
|---|---|---|---|---|---|
| 11 | spectral | cor | 0.2929 | 0.23018 | 1.83174 |
| 12 | hier | jaccard | 0.68264 | 0.40973 | 1.02991 |
| 12 | spectral | jaccard | 0.58612 | 0.19653 | 2.45999 |
| 12 | hier | cosine | 0.3507 | 0.33164 | 1.25203 |
| 12 | spectral | cosine | 0.29515 | 0.28504 | 1.85932 |
| 12 | hier | cor | 0.49478 | 0.38716 | 1.08105 |
| 12 | spectral | cor | 0.45381 | 0.31444 | 1.8129 |
| 13 | markov | jaccard | 0.48748 | 0.59323 | 1.04356 |
| 13 | hier | jaccard | 0.45007 | 0.39865 | 0.95092 |
| 13 | spectral | jaccard | 0.45896 | 0.40584 | 1.64412 |
| 13 | markov | cosine | 0.43511 | 0.65671 | 1.15911 |
| 13 | hier | cosine | 0.52333 | 0.36476 | 1.06308 |
| 13 | spectral | cosine | 0.27627 | 0.38293 | 1.57875 |
| 13 | markov | cor | 0.27984 | 0.59014 | 0.94866 |
| 13 | hier | cor | 0.5155 | 0.43232 | 0.95523 |
| 13 | spectral | cor | 0.40172 | 0.44331 | 1.42728 |
| 14 | markov | jaccard | 0.4076 | 0.52711 | 0.90023 |
| 14 | hier | jaccard | 0 | 0.3425 | 1.25433 |
| 14 | markov | cosine | 0 | 0.36638 | 1.23888 |
| 14 | hier | cosine | 0 | 0.34328 | 1.19179 |
| 14 | markov | cor | 0 | 0.48436 | 1.02836 |
| 14 | hier | cor | 0 | 0.39153 | 1.12954 |
| 15 | markov | jaccard | 0.48478 | 0.65917 | 0.6688 |
| 15 | hier | jaccard | 0.42981 | 0.54755 | 0.86747 |
| 15 | spectral | jaccard | 0.25825 | 0.37461 | 1.57243 |
| 15 | markov | cosine | 0.38534 | 0.65662 | 0.63388 |
| 15 | hier | cosine | 0.52594 | 0.48425 | 0.88447 |
| 15 | spectral | cosine | 0 | | 1.18501 |
| 15 | markov | cor | 0.38009 | 0.72949 | 0.60987 |
| 15 | hier | cor | 0.56565 | 0.50228 | 0.77945 |
| 15 | spectral | cor | 0.025 | 0.42262 | 1.25565 |
| 16 | markov | jaccard | 0.81146 | 0.64545 | 0.62651 |
| 16 | hier | jaccard | 0.62081 | 0.37608 | 0.96969 |
| 16 | spectral | jaccard | 0.45826 | 0.07458 | 3.15218 |

| | | | | | |
|---|---|---|---|---|---|
| 16 | markov | cosine | 0.66612 | 0.54656 | 0.77476 |
| 16 | hier | cosine | 0.42265 | 0.31512 | 1.08316 |
| 16 | spectral | cosine | 0.4906 | 0.10553 | 2.7723 |
| 16 | markov | cor | 0.61023 | 0.5214 | 0.7755 |
| 16 | hier | cor | 0.41574 | 0.33522 | 1.07019 |
| 16 | spectral | cor | 0.24896 | 0.12284 | 2.65623 |
| 17 | markov | jaccard | 0.48483 | 0.54416 | 0.82846 |
| 17 | hier | jaccard | 0.48955 | 0.34784 | 1.09449 |
| 17 | spectral | jaccard | 0.24658 | 0.38211 | 1.54395 |
| 17 | markov | cosine | 0.46297 | 0.49542 | 0.98161 |
| 17 | hier | cosine | 0.30804 | 0.34553 | 1.19259 |
| 17 | spectral | cosine | 0.16987 | 0.54673 | 1.21856 |
| 17 | markov | cor | 0.40352 | 0.57238 | 0.82113 |
| 17 | hier | cor | 0.46478 | 0.45088 | 0.94292 |
| 17 | spectral | cor | 0.04874 | 0.52076 | 1.27399 |
| 18 | markov | jaccard | 0.68826 | 0.67304 | 0.57419 |
| 18 | hier | jaccard | 0.64436 | 0.53479 | 0.76654 |
| 18 | spectral | jaccard | 0.3061 | 0.20473 | 2.58228 |
| 18 | markov | cosine | 0.5873 | 0.63353 | 0.56437 |
| 18 | hier | cosine | 0.53535 | 0.55286 | 0.71937 |
| 18 | spectral | cosine | 0.10484 | 0.33363 | 1.79958 |
| 18 | markov | cor | 0.69589 | 0.67661 | 0.53762 |
| 18 | hier | cor | 0.52122 | 0.58346 | 0.68757 |
| 18 | spectral | cor | 0.10195 | 0.30864 | 2.05184 |
| 19 | markov | jaccard | 0.53056 | 0.66292 | 0.66523 |
| 19 | hier | jaccard | 0.54468 | 0.47598 | 0.90583 |
| 19 | spectral | jaccard | 0.34256 | 0.38996 | 1.47986 |
| 19 | markov | cosine | 0.17374 | 0.44669 | 1.01181 |
| 19 | hier | cosine | 0.44287 | 0.42369 | 0.95953 |
| 19 | spectral | cosine | 0.15754 | 0.50891 | 1.17335 |
| 19 | markov | cor | 0.36075 | 0.5782 | 0.73762 |
| 19 | hier | cor | 0.42173 | 0.4854 | 0.84102 |
| 19 | spectral | cor | 0.2324 | 0.51015 | 1.2916 |
| 20 | markov | jaccard | 0.70568 | 0.55863 | 0.67938 |

| | | | | | |
|---|---|---|---|---|---|
| 20 | hier | jaccard | 0.69872 | 0.40465 | 0.89001 |
| 20 | spectral | jaccard | 0.32739 | 0.0453 | 4.08377 |
| 20 | markov | cosine | 0.67079 | 0.58796 | 0.75775 |
| 20 | hier | cosine | 0.51805 | 0.41542 | 0.91892 |
| 20 | spectral | cosine | 0.07012 | 0.09667 | 2.59817 |
| 20 | markov | cor | 0.65258 | 0.62181 | 0.70445 |
| 20 | hier | cor | 0.45168 | 0.45389 | 0.83463 |
| 20 | spectral | cor | 0.35472 | 0.22621 | 2.10047 |

**Supplementary Methods**

Generating the input for *aPEAR*, *emapplot* and *Enrichment Map* visualisations

To generate the enrichment networks using the *emapplot* (Yu 2022) and the *Enrichment Map* (Merico *et al.* 2010), we performed gene set enrichment analysis using the example gene list from R package *DOSE* (Yu *et al.* 2015). As *emapplot* works best with *gseaResult* (or *enrichResult*) objects as input, we performed the analysis using the *gseGO* method from *clusterProfiler* (Wu *et al.* 2021) with default parameters and cellular component gene ontology. For the *Enrichment Map* we performed the analysis using the GSEA software (Mootha *et al.* 2003; Subramanian *et al.* 2005) with 1000 permutations, classic scoring scheme and *meandiv* normalisation method; the pathways were filtered to contain a minimum of 10 and a maximum of 200 genes. We used the GMT file provided by the Bader Lab, Toronto (Human GO:BP All Pathways).

*Cytoscape* and *Enrichment Map*

The Cytoscape software and its plugin *Enrichment Map* (Merico *et al.* 2010) can be used to generate an enrichment network. However, this process is difficult to automate, requires manual rerun when the enrichment results change, and the visualisation may still require some improvements by hand. We used the R package *RCy3* (Gustavsen *et al.* 2019) to generate the enrichment network using the Cytoscape software and its plugin *Enrichment Map* directly from an R script. It requires that the Cytoscape software is running in the background, also, we had to include a sleeping time period in order to make sure that the results in the Cytoscape software are properly updated. We used the enrichment results obtained from the GSEA software and ran the *enrichmentmap mastermap* command with q-value cutoff 0.05, similarity cutoff 0.85 and did not perform filtering by expression. Then, we removed the edges and nodes with few connections and annotated the graph with *autoannotate* (Kucera *et al.* 2016).

*emapplot*

The *emapplot* visualisation was created using the *clusterProfiler (Wu et al. 2021)* enrichment output. First, pairwise-termism was calculated using the *pairwise_termsim* method from the R package *enrichplot* (Yu 2022), with the *showCategory* parameter set to the size of the complete enrichment dataset. Then, the *emapplot* function was used to generate the enrichment network with node colour set to *NES* and node label set to *group*.

Cluster quality evaluation

We used 10 different real-life datasets (Supplementary Table 1) and performed enrichment analysis using *clusterProfiler* (Wu *et al.* 2021), as well as the GSEA software (Mootha *et al.* 2003; Subramanian *et al.* 2005). We used the GMT file provided by the Bader Lab, Toronto (Human GO:BP All Pathways). This resulted in 20 different gene sets which were then used to evaluate the clustering quality with all the implemented similarity metrics (Jaccard index, cosine similarity score, and correlation) and clustering algorithms (hierarchical, spectral, and Markov clustering). In total, we measured the quality of 180 gene set clusters obtained from *aPEAR* (Supplementary Table 2).
